# Auditory-cognitive determinants of speech-in-noise perception: structural equation modelling of a large sample

**DOI:** 10.1101/2025.07.10.664090

**Authors:** Ester Benzaquén, Hasan Çolak, Xiaoxuan Guo, Meher Lad, Steven P. Rushton, Timothy D. Griffiths

**Affiliations:** Biosciences Institute, Newcastle University, Newcastle Upon Tyne, United Kingdom; Department of Audiology, Hacettepe University, Ankara, Türkiye; Translational and Clinical Research Institute, Newcastle University, Newcastle Upon Tyne, United Kingdom; School of Natural and Environmental Sciences, Newcastle University, Newcastle Upon Tyne, United Kingdom

**Keywords:** speech-in-noise, structural equation modelling, auditory working memory, auditory processing, ageing

## Abstract

Problems understanding speech-in-noise (SIN) are commonly associated with peripheral hearing loss. But pure tone audiometry (PTA) alone fails to fully explain SIN ability. This is because SIN perception is based on complex interactions between peripheral hearing, central auditory processing (CAP) and other cognitive abilities. We assessed interaction between these factors and age using a multivariate approach that allows the modelling of directional effects on theoretical constructs: structural equation modelling. We created a model to explain SIN using latent constructs for sound segregation, auditory (working) memory, and SIN perception, as well as PTA, age and measures of non-verbal reasoning. In a sample of 207 participants aged 18-81 years old, age was the biggest determinant of SIN ability, followed by auditory memory. PTA did not contribute to SIN directly, although it modified sound segregation ability, which covaried with auditory memory. A second model, using a CAP latent structure formed by measures of sound segregation, auditory memory, and temporal processing, revealed CAP to be the largest determinant of SIN ahead of age. Furthermore, we demonstrated the impact of PTA and non-verbal reasoning on SIN are mediated by their influence on CAP. Our results highlight the importance of central auditory processing in speech-in-noise perception.

## 1. Introduction

Approximately 1 in 4 people, over 1.3 billion worldwide, have hearing loss, and those of advanced age are overly impacted (Akeroyd & Munro, 2024; Vos et al., 2016). With an ageing population, hearing loss poses an increasing societal problem. Older adults suffering from hearing loss not only face a world without sound, but experience impaired communication, social isolation, and increased risk of depression (Li et al., 2014; Mener et al., 2013) and dementia (Gallacher et al., 2012; Griffiths et al., 2020; Lin et al., 2011; Lin et al., 2013; Livingston et al., 2024)

Older adults who struggle to hear in everyday life situations often complain about their inability to understand speech, especially in adverse conditions, such as with competing speakers in the background, environmental noises, or low or degraded speech (Kochkin, 2010). Speech-in-noise (SIN) perception, colloquially termed the “cocktail party problem”, has been the target of hearing research for several decades. Although a person’s overall hearing ability, measured by pure tone audiometry (PTA), is a major factor determining speech-in-noise perception, it fails to fully explain it (Anderson et al., 2013; Füllgrabe et al., 2015; Griffiths, 2023). Additionally, the use of hearing aids is unable to fully resolve SIN difficulties, as demonstrated by the dissatisfaction of some hearing aid users in noisy situations and in large group settings (Kochkin, 2000, 2005).

One reason that PTA cannot fully explain speech-in-noise ability nor interventions that only target sound levels (i.e., hearing aids) can fully restore it, is because SIN perception is inherently more complex than a process solely reliant on peripheral auditory mechanisms; it also requires central sound processing, involving the brainstem and cortex (Ding & Simon, 2012; Chandrasekaran & Kraus, 2010). When speech embedded in noise reaches a person’s cochlea, the target speech still needs to be isolated from the background and remembered until meaning can be extracted. This complex process demands both sound segregation ability and working memory capacity.

Sound segregation has been previously studied using artificial stimuli that simulate actual speech in noise. In these tasks, participants hear a complex auditory stimulus within which a target sound has to be detected. We developed a Stochastic Figure Ground (SFG) paradigm, which aims to emulate the necessity of extracting a meaningful signal from a perceptually similar background but with no linguistic information (Teki et al., 2013; Teki et al., 2011). SFG is comprised of pure-tone components that repeat over time to form a figure (target) while random tones varying over time form the ground (masker). Previous research has been successful at linking SFG and speech-in-noise ability, as well as proving the involvement of higher-level brain structures beyond the auditory cortex in its processing (Guo, Benzaquén, et al., 2025; Holmes & Griffiths, 2019; Holmes et al., 2021; Teki et al., 2016; Teki et al., 2011). Further, SFG can elicit similar neural entrainment to that of speech (Guo, Mai, et al., 2025; O’Sullivan et al., 2015).

Working memory capacity has been repeatedly linked to speech-in-noise perception (for reviews, see Akeroyd, 2008; and Dryden et al., 2017). From first principles, understanding speech in adverse conditions when the perceptual signal is not clear, such as in speech in noise paradigms, requires a level of “post-processing” of the input sound to decode it, for example, to support postdiction - the reconstruction of misheard words. Thus, it is not surprising that SIN relies partly on our ability to maintain and manipulate sounds in mind. For example, working memory is associated with noise-vocoded speech perception (Rosemann et al., 2017) and with the processing of speech from competing sources (James et al., 2014) – both adverse listening conditions similar to SIN. Working memory may even be the mechanism behind better SIN in musicians (Anderson et al., 2013; Lad et al., 2022). It has been proposed (Fullgrabe & Rosen, 2016) that working memory is especially important when the periphery starts to degrade -e.g., due to age- and the sound input lacks fidelity. In this view, working memory acts to compensate for inaccurate perception. Nevertheless, short-term auditory memory must be, axiomatically, involved in sentence-level speech comprehension; a sentence being processed always needs to be retained *in mind* until this sentence is completed (or can be accurately predicted) before it can be understood.

Speech-in-noise perception relies on other cognitive abilities beyond central auditory processing, for example, processing speed, inhibitory control, and crystallized intelligence (Akeroyd, 2008; Dryden et al., 2017). It is generally understood that as hearing deteriorates, speech perception requires greater input from cognition (Wayne & Johnsrude, 2015). This may extend to speech-in-noise perception, where the sensory input is also “deteriorated”. For example, nonverbal reasoning is associated with the recognition of degraded speech in both cochlear implant users and normal-hearing adults (Mattingly et al., 2018). Additionally, speech-in-noise perception requires less sensory evidence and relies more on preparatory (cognitive) processes (Vaden et al., 2022).

Age is undoubtedly linked to SIN ability, as peripheral hearing, central auditory processing, and cognition decrease with age (for a review, see Windle et al., 2023). Age is also known to degrade temporal processing (Anderson & Karawani, 2020), another central mechanism implicated in speech-in-noise perception (Füllgrabe et al., 2015). In older adults with and without hearing loss, cognitive function appears highly influential in SIN perception, although cognitive function is tightly related to age itself (Marsja et al., 2022). The inter-relationship amongst age, hearing, central auditory processing, and cognition complicates the interpretation of these associations. Mainly, if age leads to greater hearing loss and greater cognitive decline, as well as worse auditory processing, how can we disentangle the relationship between these variables and speech-in-noise perception? Additionally, the discrete contributions of working memory and sound segregation to speech-in-noise perception remain to be elucidated.

Here, we have taken a multivariate approach that allows us to study theoretical constructs and their complex casual relationships: structural equation modelling (SEM). SEM is a statistical method where latent constructs can be defined using measured variables or indicators, and directional links modelled based on empirical hypothesis. By creating separate latent variables for central auditory processes, as well as general intelligence as measured by a matrices test, their discrete contributions to speech-in-noise perception for both words and sentences can be assessed. Furthermore, by adding age, PTA and the causal link between them, a more accurate and detailed view of the effects of age and hearing on SIN can be discovered. We first test a structural equation model where sound segregation and auditory (working) memory are separated in two latent variables. The sound segregation construct is measured using SFG tasks, while the auditory memory construct is measured using the backwards digit span test and tests of precision for delay-matching sounds based on frequency and amplitude modulation rate (Lad et al., 2022). Considering sound segregation and auditory memory are part of an overlying theoretical construct, i.e. central auditory processing (CAP), and both rely on similar brain architecture, including primary and non-primary auditory cortex (Kumar et al., 2021; Kumar et al., 2016; Teki et al., 2016), a second structural equation model is created. In this model, sound segregation and auditory memory, in addition to temporal processing, measured with a between-channels gap detection task, are joined in a latent construct representing CAP.

## 2. Materials and Methods

### 2.1. Participants

Data from 223 participants (150 females) aged between 18-81 years old was collected. Inclusion criteria were native English-speaker status, the absence of any hearing complaints (such as self-perceived or diagnosed hearing loss, the use of hearing aids, or tinnitus), no history of neuropsychological disorders, and no current use of neurotropic medication. A total of 13 participants were removed from all analyses due to dyslexia diagnosis (2), inability to perform the sentence-in-noise task (2), non-native English-speaker status (1), and due to being duplicated data (8). For those who were tested twice (i.e., duplicated) only the earliest session was used for analysis. Thus, data from 210 participants (140 female) was used for analysis. Participants’ age ranged from 18 to 81 years (mean: 49.09; median: 51.08; standard deviation [SD]: 16.12). Data from 3 participants was incomplete and thus were not included in the SEM analysis. The study was approved by Newcastle University’s Ethics Committee (Reference numbers: 10356/2018 and 46225/2023), and written informed consent was obtained from all participants before the start of the study. The study was performed in accordance with the Declaration of Helsinki (World Medical Association, 2024).

### 2.2. Materials

#### 2.2.1. Speech-in-noise

Speech-in-noise tasks for both words and sentences were included. The word-in-noise (win) test consisted of the Iowa Test of Consonant Perception - British version (ITCP-B) (Guo et al., 2024). The ITCP-B is a single-word closed-set computer task with phonetically-balanced features. The test consists of 120 consonant-vowel-consonant words spoken by either a male or female speaker amongst 8-talker babble noise. Participants hear the target word (e.g., “moon”), which has a one second post-babble onset, and are then presented with a self-paced 4-alternative forced-choice screen with phonetically similar words (or minimal pairs) (e.g., “moon-boon-dune-noon”). Participants make their selection using the numbers 1-4 on the keyboard and are then presented with feedback (“Correct-Incorrect”) for 0.6 seconds. A new trial starts one second afterwards while a fixation cross is shown on the screen. Words are always presented at a -2 dB signal-to-noise ratio (SNR). The babble was formed by 4 female and 4 male British speakers, and the recording lasted 15 seconds. A list of pre-defined starting points for the babble was created spanning every 0.1 seconds starting at 0, and permuted without replacement for each participant. Half of the words were spoken by a male speaker while the other half by a female speaker, and this was randomly selected per participant. Participants were given a break after every 40 trials. The performance was calculated as the proportion of correct answers.

The sentence-in-noise test consisted of the British version of the Oldenburg sentences (Holmes & Griffiths, 2019). This is a closed-set test where sentences follow the structure <NAME-verb-number-adjective-noun> (e.g., “William sees four white houses”) and are spoken by a male speaker masked by 16-talker babble noise. A 5×10 matrix of all possible word combinations is presented to participants, who have to respond using a mouse. Answers are considered correct only if all selected words are correct. This is an adaptive paradigm with a 1-up 1-down staircase. The starting SNR was 10 dB, which changed in steps of 3 dB, reducing to 2 dB after the first reversal, and further reducing to 1 and 0.5 dB after 4 and 6 reversals, respectively. The babble was presented for 3.3 seconds, and sentence onset was 0.25 seconds post-babble. The masker consisted of a recording of 21.49 seconds, and its starting point is fully randomised per trial from 0 to 18 seconds. Sentence presentation is also fully randomised per trial. The task ends after 12 reversals, and the performance threshold is calculated as the median SNR of the last 6 reversals.

#### 2.2.2. SFG

Two SFG tasks were included. The SFG-Gap discrimination task was adapted from Holmes and Griffiths (2019). SFG is formed by tone chords lasting 50 ms. SFG can be divided into two components called “Figure” (target) and “Ground” (masker). The ground was composed of between 5-15 tone elements per chord, for a total of 70 chords (3.5 seconds). Tones were selected from a frequency space between 179.73 - 7246.29 Hz in a logarithmic scale. The figure was composed of 3 tone elements per chord which repeated over time for a total of 42 chords (2.1 seconds), and it started between chords 16-20. A set of 144 figures was created in advance and the order randomised to present to each participant. When necessary, a new iteration of this set was presented. Two SFG stimuli were presented per trial with an inter-stimulus interval (ISI) of 400 ms. One of the stimuli had a 6-chords-long gap in the Figure, constrained to start between chords 11-32 of the figure. Participants were required to respond to which stimulus had a gap in the figure. This is a 1-up 1-down adaptive procedure, where the target-to-masker ratio (TMR) is changed. Starting TMR was 10 dB, and TMR is changed in steps of 4 dB, which are reduced to 2 dB after the first reversal, and further reduced to 1 dB after 4 reversals. The task ended after 10 reversals. Participants were familiarised with the stimuli at the beginning of each task by introducing the concepts of “figure” and “ground”, and allowing a practice run of 6 trials at the starting TMR.

The SFG-Figure discrimination task followed the same trial structure and adaptive procedure as the SFG-Gap, but consisted of stimuli of 2 second-duration. One of the stimuli was background only while the other one included a figure spanning 6 chords, which again could start at chords 16-20, and the participants’ task was to select which stimulus contained the figure. Due to the adaptive nature of the paradigm and to avoid changes in overall power between both stimuli, the background-only stimulus included a “dummy” figure of the same duration created of random elements not already contained within the ground that changed in TMR in the same fashion. More specifically, three tone elements were added to the 6 adjacent chords representing the figure. The frequencies of said tone elements were selected randomly for each of the 6 chords. The current trial TMR was then applied to this “dummy” figure. This was done to prevent successful task completion by perceiving changes in overall power between stimuli with and without a figure. Thresholds for both tasks were calculated as the median TMR of the last 6 reversals.

#### 2.2.3. Auditory working memory

An auditory memory task to calculate memory precision for frequency (Freq) and amplitude modulation (AM) rate was used (Lad et al., 2022; Lad et al., 2024). Participants heard either a one-second pure-tone or AM-modulated white noise. After a delay of 2 seconds, the target sound had to be matched by clicking on an unlabelled and continuous visual horizontal scale representing the frequency (440 – 880 Hz) or AM rate (5 – 20 Hz) space. Frequencies were selected from a uniform distribution and a sinusoidal function was used to apply the amplitude modulation. Every click with a mouse played their selected sound, which could be done repeatedly and without time limit, after which they would press the ‘Enter’ key to confirm. The stimulus type was alternated trial-by-trial for a total of 32 trials. A break was given to participants after 16 trials. Four practice trials, 2 for each stimulus type, were presented to the participant at the beginning of the task. A precision score was obtained for frequency and AM performance by using the inverse of the standard deviation of the errors calculated using a Gaussian function.

A measure of phonological working memory was also included: the digit span (DS) test from WMS-III (Wechsler Memory Scale – Third Edition. The Psychological Corporation). Participants are required to repeat a sequence of digits increasing in load as they heard them (DS Forward) or in the opposite order (DS Backward). The total score represents the number of sequences repeated accurately.

#### 2.2.4. Between channels gap discrimination

A between-channels gap (B-C Gap) discrimination task was adapted from Phillips et al. (1997). This test was designed to be a ‘central’ gap-detection task that requires the recognition of a gap between frequencies that are represented separately in the ascending auditory pathway. Two narrow-band noises with a bandwidth of 0.25 octaves and a 0.5 ms ramp were separated by a silent interval. The first sound was centred at 4 kHz and lasted 10 ms, while the second sound was centred at 1 kHz and lasted 300 ms. The gap duration started at 200 ms and changed following a 1-up 2-down staircase paradigm, starting with a step-change of 20 ms, followed by 15 (after 3 reversals), 10 (after 6 reversals), 5 (after 8 reversals), 2 (after 10 reversals), and 1 (after 12 reversals) ms. The task ended after a total of 19 reversals or after reaching 125 trials, whichever happened first. Participants were presented with two pairs of stimuli separated by 600 ms, one with a gap as described above and one without (1 ms gap). Participants had to press (self-paced) the number keys ‘1’ or ‘2’ depending on whether the gap was in the first or second position. Feedback was shown on the screen (‘Correct!’ or ‘Wrong!’) for 500 ms. A new trial (inter-trial interval) started after 1 second. During the task, if the gap duration reached 1 ms, any answer was considered wrong and the gap duration increased. Before the beginning of the task, and after a familiarisation run where participants were introduced to the target stimuli, 12 practice trials were presented where the gap duration was 230 ms, and it adaptively changed in a 1-up 1-down pattern by 5 ms. The performance threshold was calculated as the median duration of the gap on the last 6 reversals.

#### 2.2.1. General (fluid) Intelligence

A matrix test to measure general or fluid intelligence was created using the matrix reasoning item bank (MaRs-IB; Chierchia et al., 2019). Matrices were all taken from set 1. Participants familiarised themselves with the task with 4 practice trials using the first matrices from the set (numbers 1 – 4). The test included a total of 26 matrices; 25 in sequential order starting with item 6 (numbers 6 – 30), and the matrix number 47 of greater difficulty to avoid ceiling effects. Participants had 30 seconds to respond to each matrix, and a countdown timer appeared for the last 5 seconds. A total score representing the number of correct matrices was created (0 – 26) per participant.

#### 2.2.6. Other measures

Measures for musicality (Goldsmith Musical Sophistication Index; Gold-MSI), premorbid intelligence and literacy (Wechsler Test of Adult Reading; WTAR), and self-reported SiN ability (Spatial Speech Questionnaire, SSQ) were also taken. These measures are not included in the current analyses and thus are not described further.

### 2.3. Procedure

After arriving at the lab and providing informed consent, PTA thresholds were measured for frequencies 0.25 – 8 kHz in a soundproof room using air conduction only with the diagnostic audiometer AD226 by Interacoustics. The computer tasks were then performed in the same soundproof room in the following order: SIN tests (words, then sentences), SFG-Figure discrimination, auditory memory (Freq + AM), SFG-Gap discrimination, Matrices, Gold-MSI, and Gap detection. Paper tests were then completed in another room in the following order: DS Forward, DS Backward, WTAR, and SSQ. The testing session usually lasted 2 hours and participants received compensation for their time. Most computer tasks were coded in JavaScript and run using Chrome, except WiN and Gap detection, which were coded using Matlab R2017a. All stimuli were presented at 65-73 dB SPL (sound pressure level).

### 2.4. Analysis

Before SEM analyses, data were linearly transformed to reduce differences in variance amongst variables and so that positive values would reflect better performance. Thus, the scores of SiN, both SFG tasks, and B-C Gap were inverted. SiN was further multiplied 10 while B-C Gap was multiplied by 0.1. WiN was changed from reflecting proportion to reflecting percentage. Lastly, the precision scores for AM and Freq were multiplied by 50 and 10, respectively.

Structural equation models (SEM) were built using the lavaan package (version 0.6-17) in R (version 4.2.1). To calculate the models, maximum likelihood estimation with nonnormality correction based on the Satorra-Bentler scaled test statistic was used (Brosseau-Liard & Savalei, 2014; Brosseau-Liard et al., 2012). The α level used for significance testing of path coefficients was set at 5%. The models were evaluated by a set of criteria using several goodness-of-fit measures: the Bentler comparative fit index (CFI), Tucker-Lewis Index (TLI), the root-mean-square error of approximation (RMSEA), and the standardised root mean squared residual (SRMR) (Hu & Bentler, 1999; Kline, 2011). Only robust measures of these indices are reported in this study (Brosseau-Liard & Savalei, 2014; Brosseau-Liard et al., 2012). Bootstrapped 95% confidence intervals were created for each model fit measure using 1000 repetitions with the ‘Bollen-Stine’ method (Bollen & Stine, 1992) as implemented in lavaan.

The choice of scaling variables for each latent construct was decided on theory alone. Due to previous research showing good predictability of SIN with SFG-Gap (Holmes & Griffiths, 2019), this was selected as the scaling variable of the SFG latent construct. For auditory working memory, recent findings implicate memory for amplitude modulation rates as one of the greatest factors determining speech-in-noise perception (Lad et al., 2024), thus AM was used as the scaling variable for the AWM construct. Because word-in-noise perception forms the basis for sentence-in-noise perception, this (i.e., win) was used as the scaling variable for SIN. For a Central Auditory Processing (CAP) latent variable, the working memory component was used as the scaling variable, as the contribution of working memory is the most replicated finding in speech-in-noise research (Dryden et al., 2017). Although scaling variables are not estimated nor tested for significance, as lavaan uses the fixed-marker technique, these are assumed significant for visualization purposes only.

## 3. Results

### 3.1. Hearing Thresholds

Participants averaged thresholds over all frequencies (0.25-8kHz) and both ears were 11.887 (± 9.294) dB hearing level (HL). No participant had greater than mild-hearing loss (<40 dB) when averaged over all frequencies across both ears. Individual and average thresholds are plotted in Figure 1.

**Figure 1.**
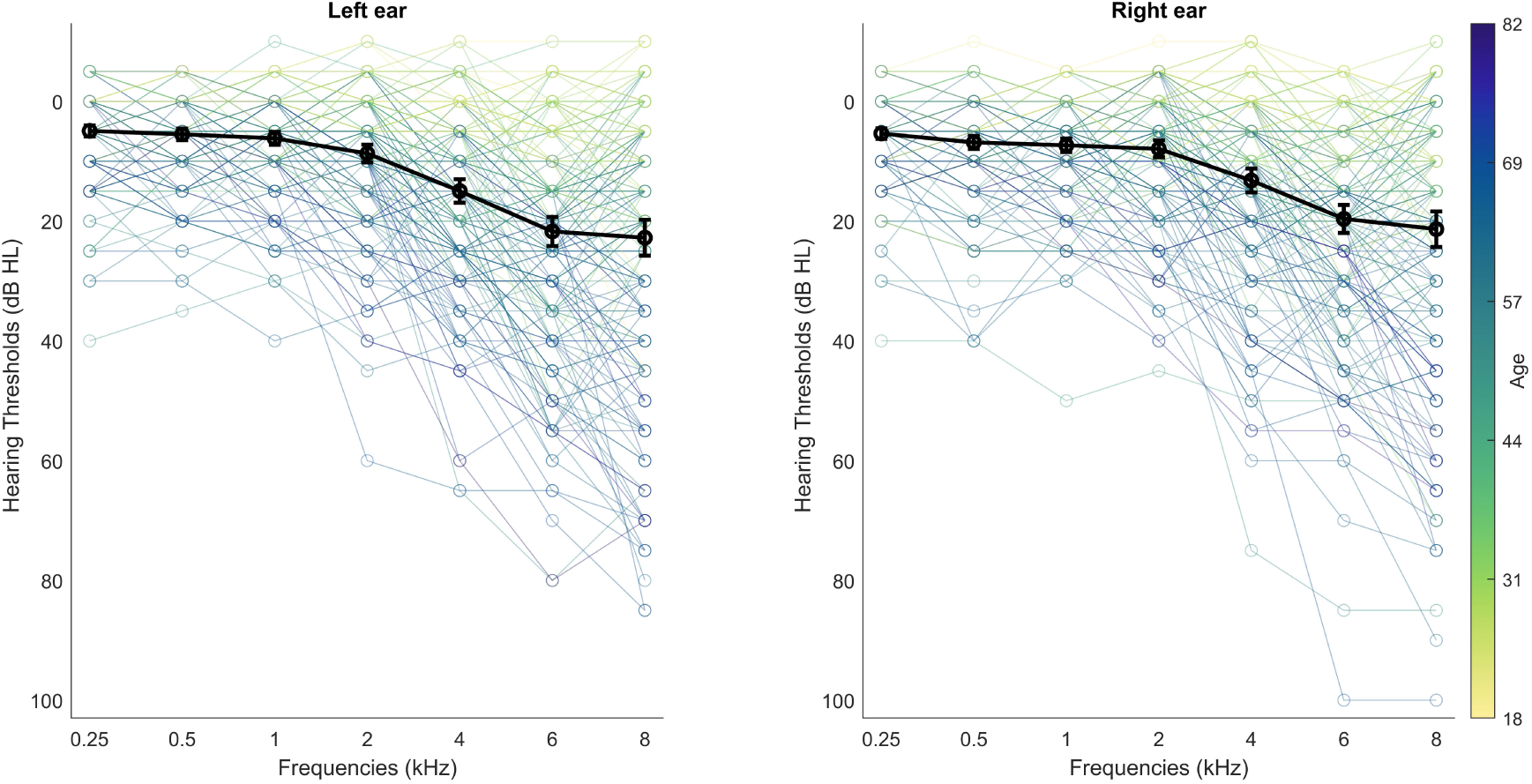
Pure tone Audiogram. Hearing thresholds in dB HL for the left and right ear. Individual data is colour-coded based on the age of the participant. Average thresholds are plotted in black.

### 3.2. Demographic data and performance

Demographic data and the average performance of all tasks are shown in Table 1. A Spearman correlation matrix was performed using Holm-Bonferroni correction for multiple comparisons. Most variables were correlated to each other, although age did not correlate with DS backwards. DS backwards also showed no correlation to PTA or both SFG tasks. Lastly, between-channels gap detection did not correlate with SFG for Figure discrimination.

**Table 1.** Descriptive statistics and Spearman correlation coefficients. Significant correlations after Holmes-Bonferroni correction are highlighted in bold.

|  | Mean (SD) | 2 | 3 | 4 | 5 | 6 | 7 | 8 | 9 | 10 | 11 |
| --- | --- | --- | --- | --- | --- | --- | --- | --- | --- | --- | --- |
| 1. Age | 49.23 (16.02) | **0.79** | **-0.72** | **0.63** | 0.01 | **-0.37** | **-0.25** | **0.32** | **0.54** | **0.31** | **-0.37** |
| 2. PTA | 12.12 (9.41) |  | **-0.68** | **0.59** | -0.06 | **-0.41** | **-0.29** | **0.27** | **0.57** | **0.36** | **-0.37** |
| 3. WiN | 0.67 (0.09) |  |  | **-0.66** | **0.20** | **0.45** | **0.42** | **-0.33** | **-0.56** | **-0.45** | **0.42** |
| 4. SiN | -2.22 (1.69) |  |  |  | **-0.20** | **-0.36** | **-0.33** | **0.23** | **0.48** | **0.33** | **-0.33** |
| 5. DS-bck | 7.29 (2.15) |  |  |  |  | **0.33** | **0.33** | -0.03 | -0.13 | **-0.29** | **0.25** |
| 6. AM | 0.22 (2.10) |  |  |  |  |  | **0.57** | **-0.27** | **-0.44** | **-0.53** | **0.42** |
| 7. Freq | 0.09 (1.53) |  |  |  |  |  |  | **-0.21** | **-0.36** | **-0.49** | **0.48** |
| 8. SFG-F | -6.65 (4.69) |  |  |  |  |  |  |  | **0.40** | 0.21 | **-0.23** |
| 9. SFG-G | -23.14 (8.45) |  |  |  |  |  |  |  |  | **0.38** | **-0.31** |
| 10. B-C Gap | 79.95 (96.57) |  |  |  |  |  |  |  |  |  | **-0.39** |
| 11. MTX | 14.24 (4.42) |  |  |  |  |  |  |  |  |  |  |

### 3.3. Sound Segregation and Auditory (Working) Memory as separate contributors to speech-in-noise

A SEM model was constructed with separate latent variables representing sound segregation (‘SFG’) and Auditory Working Memory (‘AWM’). The contributions of age, PTA, and overall intelligence (‘MTX’) were also included in the model. This model (Model 1), including path coefficients and model fit indices, can be seen in Figure 2.

**Figure 2.**
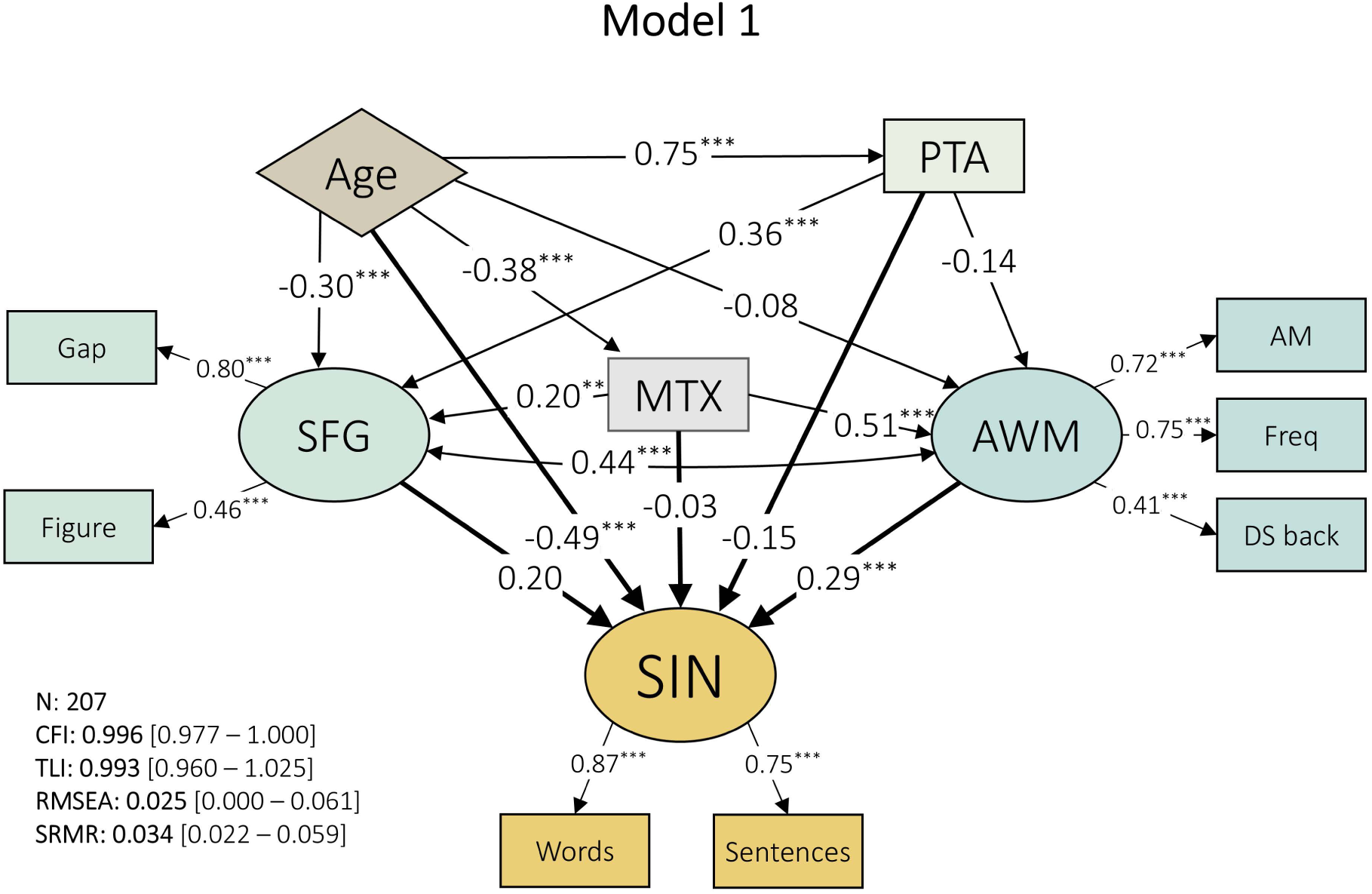
Model 1. Path coefficients are shown within the arrows representing the paths. Latent variables are represented with elliptic shapes, while indicators and observed variables are denoted with rectangles. The exogenous variable is plotted in a diamond shape. Model fit indices on the bottom left corner with bootstrapped 95% confidence intervals in square brackets. * = p<0.05, ** = p< 0.01, *** = p<0.001.

Model 1 (Figure 2) had a good model fit as demonstrated by the CFI (0.996, 95% CI: 0.977-1) and RMSEA (0.025, 95% CI: 0-0.061). This model was able to explain 83.172% of the variance of speech-in-noise (R^2^: 0.832, Adjusted R^2^: 0.828). Based on path coefficients (β), the factor that had the greatest effect on speech-in-noise was age (β: -0.49, p<0.001), followed by Auditory Working Memory (β: 0.29, p<0.01). Further, age significantly modified hearing (PTA; β: 0.75, p<0.001), sound segregation (SFG; β: -0.3, p<0.001), and general intelligence (MTX; β: -0.38, p<0.01). Although intelligence contributed to SFG (β: 0.2, p<0.01) and AWM (0.51, p<0.001), it did not have a direct effect on SIN (-0.03, p>0.05). Similarly, PTA and SFG did not predict SIN changes significantly (PTA; β: -0.15, p>0.05; SFG; β: 0.25, p>0.05). The contribution of hearing (PTA) on SIN ability was also not mediated by (its effect on) working memory (β: -0.14, p>0.05). Hearing (PTA) had only an effect on sound segregation (SFG; β: -0.36, p<0.001). The shared variance between working memory and sound segregation was large and statistically significant (β: 0.44, p<0.01).

### 3.4. The effects of Central Auditory Processing on speech-in-noise

An alternative model (Model 2; Figure 3) was created where sound segregation and working memory were joined under a general process of CAP, with one indicator each. SFG-Gap was used to exemplify sound segregation ability, while auditory memory for amplitude modulation (AM) was used to index auditory working memory. A third indicator was used to represent temporal processing: between-channels gap discrimination.

**Figure 3.**
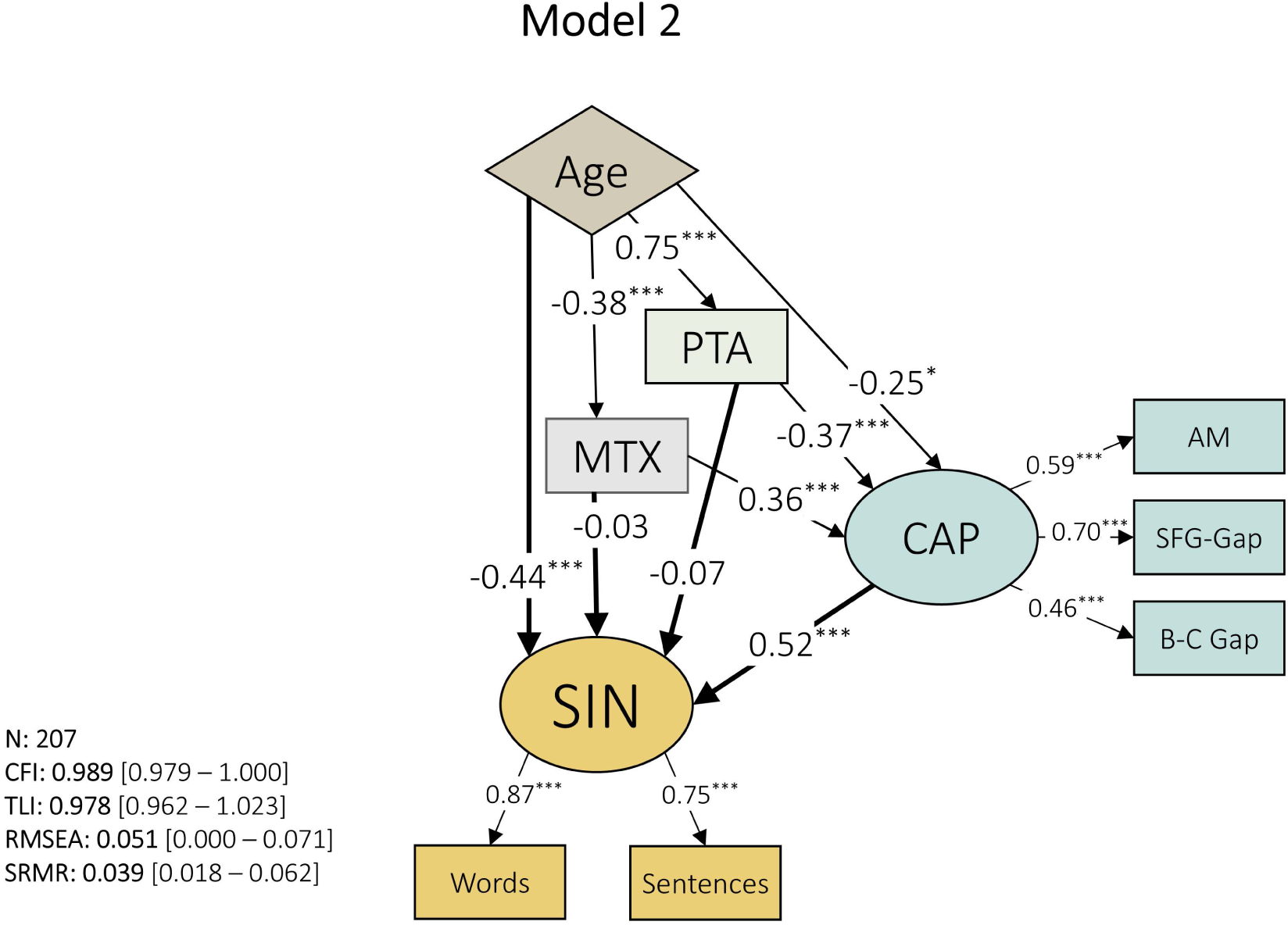
Model 2. A central auditory process (CAP) latent variable was created to include measures of sound segregation (SFG-Gap), auditory working memory (AM), and temporal processing (between-channels Gap detection). Path coefficients are shown within the arrows representing each path. Latent variables are represented with elliptic shapes, while indicators and observed variables are denoted with rectangles. The exogenous variable is plotted in a diamond shape. Model fit indices on the bottom left corner with bootstrapped 95% confidence intervals in square brackets. * = p<0.05, ** = p< 0.01, *** = p<0.001.

By combining three types of indicators into one latent construct – CAP – this factor achieved a significant contribution to SIN with a path coefficient of 0.52 (p<0.001), greater than that of age (β: -0.44, p<0.001). The relationship between SIN and PTA (β: -0.07, p>0.05), and SIN and MTX (β: -0.03, p>0.05), was mediated through their effects on CAP (PTA→CAP, β: -0.37, p<0.001; MTX→CAP, β: 0.36, p<0.001). Age had a causal effect on CAP (β: -0.25, p<0.05), PTA (β: 0.75, p<0.001), and MTX (β: -0.38, p<0.001).

Model fit, although slightly smaller than the previous model (Model 1), was good as exemplified by CFI (0.989, 95% CI: 0.979-1) and RMSEA (0.051, 95% CI: 0-0.071). This model was able to explain 83.802% of the variance of speech-in-noise (R^2^: 0.838, Adjusted R^2^: 0.835), slightly higher than the previous model despite using fewer variables.

All path coefficients for Models 1 and 2 between latent variables, including observed variables, can be seen in Table 2.

**Table 2.** Path coefficients between latent variables as well as exogenous variables for Model 1 and 2.

| Model | Variable | Regressor | $\beta$ [95% CI] | Z | p |
| --- | --- | --- | --- | --- | --- |
| Model 1 | SIN | Age | -0.485 [-0.655 -0.314] | -5.566 | <0.001 |
|  |  | PTA | -0.147 [-0.317 0.022] | -1.705 | 0.088 |
|  |  | SFG | 0.200 [-0.069 0.468] | 1.459 | 0.145 |
|  |  | AWM | 0.291 [0.099 0.482] | 2.971 | 0.003 |
|  |  | MTX | -0.026 [-0.172 0.119] | -0.351 | 0.726 |
|  | SFG | Age | -0.300 [-0.541 -0.085] | -2.732 | 0.006 |
|  |  | PTA | -0.364 [-0.573 -0.154] | -3.406 | 0.001 |
|  |  | AWM~ | 0.443 [0.201 0.685] | 3.586 | <0.001 |
|  |  | MTX | 0.201 [0.063 0.340] | 2.856 | 0.004 |
|  | AWM | Age | -0.081 [-0.292 0.130] | -0.752 | 0.452 |
|  |  | PTA | -0.139 [-0.330 0.052] | -1.427 | 0.154 |
|  |  | SFG~ | 0.443 [0.201 0.685] | 3.586 | <0.001 |
|  |  | MTX | 0.508 [0.382 0.633] | 7.908 | <0.001 |
|  | MTX | Age | -0.380 [-0.481 -0.278] | -7.338 | <0.001 |
|  | PTA | Age | 0.746 [0.699 0.793] | 31.012 | <0.001 |
| Model 2 | SIN | Age | -0.439 [-0.620 -0.257] | -4.733 | <0.001 |
|  |  | PTA | -0.069 [-0.249 0.112] | -0.744 | 0.457 |
|  |  | CAP | 0.522 [0.271 0.773] | 4.078 | <0.001 |
|  |  | MTX | -0.027 [-0.186 0.132] | -0.334 | 0.738 |
|  | CAP | Age | -0.247 [-0.450 -0.045] | -2.394 | 0.017 |
|  |  | PTA | -0.368 [-0.561 -0.174] | -3.720 | <0.001 |
|  |  | MTX | 0.362 [0.232 0.491] | 5.479 | <0.001 |
|  | MTX | Age | -0.380 [-0.481 -0.278] | -7.338 | <0.001 |

~ after regressor name represents covariance

To assess multicollinearity, which could weaken the confidence in coefficient estimates, we built a linear regression model. First, sentence and word scores were standardized and summed together to create the dependent variable, and all other measured variables used to construct both SEM (Model 1 and Model 2) were defined as predictors. Then, tolerance and variance inflation factors (VIF) were calculated using the package “olsrr” in R. VIF values > 10 are considered to indicate potential collinearity, and thus could undermine the interpretation of the models (Chatterjee & Hadi, 2006). VIF values, which can be seen in Table 3, were greatest for Age and PTA, but these were far from the multicollinearity threshold (2.50 and 2.44, respectively).

**Table 3.** VIF and Tolerance values for predictors used in SE.

|  | Tolerance | VIF |
| --- | --- | --- |
| Age | 0.400 | 2.501 |
| PTA | 0.409 | 2.442 |
| Freq | 0.606 | 1.649 |
| AM | 0.594 | 1.683 |
| SFG-Figure | 0.830 | 1.205 |
| SFG-Gap | 0.594 | 1.685 |
| DS-backward | 0.846 | 1.182 |
| B-C Gap | 0.791 | 1.264 |
| MTX | 0.676 | 1.479 |

## 4. Discussion

In the current study, we constructed a structural equation model (SEM) of speech-in-noise perception (SIN). SEM is a complex multivariate approach that allows for the exploration of causal relationships between theoretical constructs. We included the observed/exogenous variables age, hearing thresholds (PTA), and general intelligence (MTX), and created latent constructs for sound segregation (SFG) and auditory working memory (AWM). By modelling the effects of age on hearing thresholds and the influence of PTA on central auditory processing, we found that hearing did not have a direct impact on speech in noise. This is in contrast to previous research that highlighted the importance of hearing thresholds over age on speech-in-noise perception (Billings & Madsen, 2018; Lad et al., 2024). One possibility for this discrepancy may be due to the fact that the majority of people in our sample had normal or near-normal hearing, with average hearing thresholds not exceeding 40 dB. Additionally, when modelling complex relationships in normal-hearing people between auditory-cognitive factors and hearing, other research has demonstrated little or no effect of audiometric thresholds (Anderson et al., 2013). Nevertheless, the inclusion of very young people with no hearing loss and the fact that our sample has only mild-hearing loss that is highly age-related, prevent us from drawing strong conclusions on the lack of direct relationships between PTA and SIN. We are unfortunately unable to assess whether extended high-frequency (>8 kHz) hearing level would be a better predictor than standard PTA as suggested by others (Çolak et al., 2024; Motlagh Zadeh et al., 2019; Yeend et al., 2019); although this finding is not always replicated (Smith et al., 2019).

We found that age was the biggest contributor to speech-in-noise perception. The unique variance explained by age was not captured by any of our auditory-cognitive variables. Previous research has highlighted the importance of age (e.g., Besser et al., 2015), yet the non-specific impacts of ageing preclude a single interpretation. Although the influence of age on some level of cognitive decline was seen in its significant relationship to nonverbal intelligence (as measured by MTX), MTX did not show any direct impact on speech-in-noise thresholds. Due to the significant direct path between age and SIN, part of the age effects on speech-in-noise difficulties relies on mechanisms not explained by our model. One possible factor may be age-related reduced inhibition, as research has demonstrated that, for people with normal hearing, deficits in speech-in-noise perception are mediated by impaired inhibitory processes (Gomez-Alvarez et al., 2023). Research has found that older adults over-represent sound signals in the cortex, and this extends to the neural processing of “unattended” speech (Presacco et al., 2016), thus it is plausible that impaired inhibition to “noise” signals hinders speech-in-noise perception. Another factor is likely the age-related slowing in processing speed, which has been associated with SIN perception (Dryden et al., 2017; Moberly et al., 2019). However, the reduced recognition of time compressed speech, which is highly affected by ageing, does not seem associated with cognitive processing speed (Tinnemore et al., 2022), casting doubts on this hypothesis. Nevertheless, age has a multifactorial effect which includes deterioration of overall cognitive abilities (Harada et al., 2013), temporal processing (Anderson & Karawani, 2020), frequency selectivity (Regev et al., 2023), and binaural processing (Moore, 2016), all of which could impact speech-in-noise perception.

Auditory working memory had a discrete effect on SIN not explained by age or hearing thresholds. As mentioned earlier, the relationship between auditory memory and speech-in-noise is the most replicated finding in SIN research (Akeroyd, 2008; Dryden et al., 2017). Although previous research has found weak or no relationship between working memory and SIN in young adults with normal hearing (Fullgrabe & Rosen, 2016), our model demonstrated an association between auditory memory and SIN independent of age and hearing levels, as they did not have significant contributions to the constructed working memory latent variable.

Sound segregation did not contribute to SIN directly, but shared a significant amount of variance with auditory working memory. Sound segregation partly relies on peripheral mechanisms, as PTA significantly affected SFG performance. According to our model, the variance explained by SFG is shared and captured by auditory memory. Besides the shared variance with auditory memory, the addition of a new SFG stimulus (figure discrimination) may be behind the lack of discrete contribution to SIN by this sound segregation construct. Other than the previously developed SFG where participants ought to discriminate the presence of a gap within the figure (Holmes & Griffiths, 2019), and which requires the tracking of the figure over time, a newly developed SFG was used that combined this type of SFG and the original version where the presence of a figure had to be detected. However, the original detection paradigm used a fixed SNR of 0 dB, such that the target and ground are only differentiated by temporal coherence (Teki et al., 2013). In the current newly developed task, which involved figure discrimination in an adaptive paradigm with varying SNR, the figure could be detected within the background by sound-level differences alone, and did not require tracking over time.

Nonverbal reasoning greatly modulated auditory working memory. It also impacted sound segregation mechanisms albeit less strongly. General intelligence is thought to determine all cognitive-related processes; however, our model stresses that the effect of general intelligence on SIN is mediated by auditory working memory. Working memory capacity and general intelligence are highly related (Conway et al., 2003), but our model considered a directional mechanism where intelligence determines memory ability. Associations between nonverbal intelligence and degraded-speech recognition have been found before (Mattingly et al., 2018; Moberly et al., 2018), but differences in processing speed may be behind this relationship (Moberly et al., 2019). The matrix test used in the current study posed a time limit to respond, and thus is confounded by processing speed, which is also undoubtedly linked to general intelligence and working memory (Fry & Hale, 2000). The relationship between intelligence and working memory may also partly be driven by the association between intelligence and sensory performance (Troche & Rammsayer, 2009), as some of the tasks used to measure working memory were based on frequency and temporal precision.

We further created a simplified SEM where sound segregation (SFG-Gap), auditory working memory (AM rate), and temporal processing (between-channel gap detection) all act under one latent structure: central auditory processing (CAP). By joining sound segregation and working memory mechanisms, and adding temporal precision, this latent construct surpassed age as the most important predictor of speech-in-noise perception. We further revealed that the impacts of hearing and intelligence on SIN act through their influence on central auditory processing. In other words, the effects of hearing and intelligence on speech-in-noise are mediated through central auditory mechanisms: greater abstract reasoning and better hearing support improved central processing, which causes enhanced comprehension of speech-in-noise. Previous research using SEM has emphasised the importance of central auditory processing to speech in noise perception. In a sample of 120 older adults, Anderson et al. (2013) found central processing was the biggest determinant of speech in noise perception, followed by cognitive function which included auditory (and working) memory.

Overall, our research demonstrates a critical role of central auditory processing, encompassing auditory (working) memory, sound segregation, and temporal precision, in speech in noise perception. Furthermore, our models emphasize the importance of non-specific age effects that go beyond declines in hearing and reasoning abilities, and how central auditory processing mechanisms mediate the effect of hearing and cognition on speech-in-noise perception. Our results suggest that CAP processes should be assessed in clinical settings to obtain a comprehensive view of the reasons for SIN difficulties.

## 6. Author contributions

E.B, X.G. and T.G. conceptualized and designed the study. T.G. received the funding. E.B. and M.L. programmed the computer tasks. E.B., H.Ç., and X.G. collected the data. E.B., H.Ç., X.G. and S.R. analysed the data. E.B. prepared all figures. E.B. wrote the original draft. T.D., H.Ç., and X.G. revised the manuscript. All authors reviewed and approved the submitted manuscript.

